# Liver X Receptor β Controls Hepatic Stellate Cell Activation via Hedgehog Signaling

**DOI:** 10.1101/577833

**Authors:** Li Zhong, Shengjie Huang, Xuan Du, Can Cai, Youping Zhou, Wei Shen, Liang Deng, Bo Ning

**Affiliations:** Department of Gastroenterology, The second Affiliated Hospital of Chongqing Medical University, Chongqing, China; Department of Gastroenterology, The First Affiliated Hospital of Chongqing Medical University, Chongqing, China; The second hospital of Dalian Medical university

**Keywords:** hepatic fibrosis, hepatic stellate cell activation, LXRβ, Hedgehog signaling

## Abstract

Liver X receptors (LXR) α and β serve important roles in cholesterol homeostasis, anti-inflammatory processes and the activation of hepatic stellate cells (HSCs). However, the development of therapies for liver fibrosis based on LXR agonists have been hampered due to side-effects such as liver steatosis. In this study, we demonstrated that HSCs expressed high levels of LXRβ, but not LXRα, and that overexpression of LXRβ suppressed fibrosis and HSC activation in a carbon tetrachloride (CCl_4_)-induced fibrosis mouse model, without resulting in liver steatosis. Furthermore, Hedgehog (Hh)-regulated proteins, markedly increased in the CCl_4_-affected liver and mainly expressed in activated HSCs, were repressed under conditions of LXRβ overexpression. In addition, LXRβ knockout led to activation of Hh signaling and triggering of HSC activation, while overexpression of LXRβ led to the inhibition of the Hh pathway and suppression of HSC activation. These results suggest that LXRβ suppresses the activation mechanism of HSCs by inhibiting Hh signaling. In conclusion, LXRβ, by restoring the differentiation of HSCs, may be a promising therapeutic target for liver fibrosis without the adverse side-effects of LXRα activation.

## Introduction

Fibrosis of the liver, triggered by chronic liver injury, is the overproduction of fibrillar collagen and remodeling of extracellular matrix. Left untreated, hepatic fibrosis can lead to scarring of the liver, known as cirrhosis, which is associated with significant morbidity and mortality rates. Fibrosis occurs primarily due to the activation of hepatic stellate cells (HSCs), which maintain a quiescent phenotype in the normal liver and store the majority of the retinyl esters, triglycerides and cholesterol esters of the body in lipid droplets [1-2]. Under conditions of liver injury, HSCs are activated, lose their lipid content, and undergo a transformation giving rise to a fibrogenic myofibroblastic phenotype [3-5]. Activated HSCs exhibit high proliferative activity and overexpress smooth muscle α actin (Acta2) and collagen, mainly of type I (collagen I) [6,7].

HSC activation and the conversion to a myofibroblastic phenotype are associated with a marked decrease in the expression of several adipogenic transcription factors, including the liver X receptors (LXRs) [8]. LXRs, occurring as two isoforms (α and β), function as sensors of cholesterol levels, prompting reverse cholesterol transport and its excretion into bile. Previous studies have revealed certain beneficial effects of these proteins on inflammation, atherosclerosis, and diabetes [9,10], making them promising therapeutic targets for these conditions. LXRs also play an important role in HSC activation and susceptibility to liver fibrosis [11,12]. Nevertheless, activation of LXRs leads to enhanced hepatic triglyceride synthesis and may give rise to liver steatosis and hypertriglyceridemia, hampering therapeutic development based on LXR agonists [13]. However, LXRα and LXRβ exhibit distinct distributions, and certain studies have indicated that it is LXRα, predominantly expressed in hepatocytes (HCs), that specifically mediates the increase of triglyceride synthesis [14,15]. Although LXRs have already been proven to influence the activation of HSCs, the specificity of the two isoforms and their molecular mechanisms remain unresolved. The aims of this research, therefore, were to investigate the antifibrogenic role of LXRβ in the process of HSC activation *in vitro* and *in vivo*. If confirmed, LXRβ-selective agonists may be a potential therapeutic target to avoid HSC activation-associated fibrosis, simultaneously avoid undesirable LXRα-associated side-effects.

In the present study, we isolated HSCs from mice and specifically silenced LXRα or LXRβ in order to evaluate the distinct roles of the isoforms in the activation of HSCs. We discovered that HSCs predominantly express LXRβ. The silencing of LXRβ by small interfering RNA (siRNA) significantly inhibited LXR target gene expression and promoted the activation of HSCs *in vitro*. Notably, LXRα knockdown had no significant effect on the expression of LXR target genes or the activation of HSCs. Moreover, the HSCs in LXRβ-overexpressing mice were resistant to carbon tetrachloride (CCl_4_)-induced activation. Hedgehog (Hh) signaling has been revealed to be important in the promotion of HSC activation and conversion to the myofibroblastic phenotype [16,17]. In chronic liver injury, Hh signaling plays a major role in liver fibrogenesis [18], therefore, its association with the protective function of LXRβ was also investigated. The results of the present study demonstrate that LXRβ regulates the activation of HSCs and prevents CCl_4_-induced fibrosis via Hh signaling, simultaneously avoiding undesirable LXRα-associated liver steatosis side-effects. Specific activators of LXRβ may be used as potential therapeutic agents against liver fibrosis.

## Materials and Methods

### 1. Animals and diets

Male C57/BL6 mice were obtained from Chongqing Medical University. All mice were housed individually in plastic cages with free access to standard chow and water. Chronic liver injury in the mice was induced by intraperitoneal (IP) injections of a 10% CCl_4_ solution in olive oil (0.5 μl pure CCl_4_/g body weight) 2 times per week for 5 weeks. Alternatively, mice were administered IP injections of LXR agonist T0901317 (50 mg/kg body weight; dissolved in DMSO at 50 mg/ml) every 3 days. The animals in the control groups received equivalent doses of DMSO and vehicle olive oil. To determine the antifibrotic effects of LXRβ, adenovirus vector (Genechem. Shanghai, China) encoding the cDNA of murine LXRβ (Ad-LXRβ) or empty control vector (Ad-control) were injected via the tail vein (5×10 ^8^ virus particles/mouse). The animal experimental procedures were performed according to the National Institutes of Health Guidelines for the Use of Experimental Animals and approved by the Ethics Committee and the Medicine Animal Care Committee of Chongqing Medical University.

### 2. Cell isolation and culture

HCs and HSCs were isolated from adult male C57 mice as previously described [19]. Briefly, the mice livers were perfused *in situ* with collagenase IV (0.5 mg/ml, Gibco; Thermo Fisher Scientific, Inc., USA) for approximately 20 min until the liver became smooth and soft. The liver was excised and placed in a dish containing Gey’s balanced salt solution (GBSS) with 0.5 mg/ml collagenase IV and 20 μg/ml DNase I. The liver capsule was opened using forceps and the cell suspension was gently dissociated using a plastic pipette. The cell suspension was filtered through a 200-μm gauze to remove undigested tissue, and centrifuged at 50 x g for 2 min at 4°C in a 50-ml tube . HCs are main components of the resulting cell pellet, while HSCs are light and remain in the supernatant. The pelleted HCs were further resuspended in 48% Percoll and purified by centrifugation at 50 x g for 10 min. The aforementioned supernatant containing the HSCs was transferred to a new 50-ml tube and centrifuged at 500 x g for 7 min at 4°C to pellet the HSCs, which were then resuspended and lysed in GBSS with 0.5 mg/ml pronase E and 20 μg/ml DNase I for 20 min at 37°C to lyse the HSCs. The mixture was centrifuged at 500 x g for 7 min at 4°C, the cell pellet was resuspended in 12 ml 15% Optiprep diluted in GBSS, and 12 ml 11.5% Optiprep was carefully transferred onto the cell suspensions, followed by 12 ml of GBSS. Following centrifugation at 1,420 x g for 20 min, the HSCs were removed from the top of the 11.5% Optiprep layer. The viability of freshly isolated cells was assessed by trypan blue exclusion, then, cultured in DMEM with 10% FBS and maintained at 37°C in a humidified 5% CO2 atmosphere. Oil red O straining or periodic acid schiff (PAS) glucogen straining was used for morphologic detection of the isolated cells. Further, the purities of primary cells were tested using glial fibrillary acidic protein (GFAP) (quiescent HSC marker; 1:100; 16825-1-AP, Proteintech Group, China) or Acta2 (activated HSC marker; 1/100; ab7817; Abcam, Cambridge, UK) for HCSs, and cytokeratin 18 (HC marker; 1:100; 10830-1-AP, Proteintech Group) for HCs (Supplementary Fig. 1). To further confirm the effects of LXRs and Hh signaling on HSC activation, after 3 days of isolation, primary HSCs were treated with LXR agonist T0901317 (5 μM) or antagonist SR9238 (10 μM), and Hh agonist Purmorphamine (2 μM) or antagonist GDC0449 (5 μM) (MedChemExpress, Monmouth Junction, NJ, USA). The siRNA targeting mouse LXRα and LXRβ sequences, constructed by Genepharma (Shanghai, China), were transfected according to the manufacturer’s instructions. The siRNA sequences were as follows: LXRα sense, 5’-GGCAACACUUGCAUCCUUATT-3’; and antisense, 5’-UAAGGAUGCAAGUGUUGCCTT-3’; LXRβ sense, 5’-CAUCCACCAUCGAGAUCAUTT-3’; and antisense, 5’-AUGAUCUCGAUGGUGGAUGTT-3’; control scramble sense, 5’-UUCUCCGAACGUGUCACGUTT-3’; and antisense, 5’-ACGUGACACGUUCGGAGAATT-3’.

### 3. Immunohistochemistry and immunofluorescence

The mouse livers were fixed in 4% paraformaldehyde and embedded in paraffin for hematoxylin-eosin (H&E), Masson, Sirius red and immunohistochemical staining. For immunochemical staining, the paraffin-embedded sections were dewaxed with xylene, rehydrated with alcohol and heated with citrate buffer for antigen retrieval. The sections were incubated with 3% peroxide to block endogenous peroxidase and then incubated in 0.5% Triton X-100 for 15 min. Following blocking with 5% goat serum solution for 40 min at 37°C, the sections were stained with anti-Acta2 (1:400) at 4°C overnight. The sections were further processed by the application of the immunoperoxidase technique, using the Envision kit (Boster Biological Technology, China). For immunofluorescence analysis, the samples were incubated with anti-LXRα (1:100; ab3585, Abcam), anti-LXRβ (1:100; ab228867, Abcam), anti-Acta2 (1:100) and anti-nuclear transcription factor Gli family zinc finger (Gli)2 (1:100; ab26056, Abcam) overnight at 4°C, followed by incubation with fluorescein isothiocyanate-labeled goat anti-rabbit and anti-mouse secondary antibody (Alexa Fluor; Invitrogen; Thermo Fisher Scientific, Inc.). DAPI was applied to each section for 10 min to counterstain the nuclei.

### 4. Western blot analysis

Cells or liver samples were lysed in radioimmunoprecipitation assay buffer with protease inhibitors, and the concentration of the resulting protein extract was measured using the bicinchoninic acid assay. The extracted proteins were separated by SDS-PAGE (10% gel) and transferred to polyvinylidene difluoride membranes. The membranes were blocked with 5% non-fat milk and immunological detection was performed with the following primary antibodies: Acta2 (1:1,000), collagen I (1:500; 14695-1-AP; Proteintech, Rosemont, IL, USA), LXRα (1:1,000), LXRβ (1:1000), sterol regulatory element binding protein-1c (Srebp-1c; 1:1000; ab28481; Abcam), ATP binding cassette subfamily A member 1 (Abca1; 1:500, Ag24118; Proteintech), Patched (Ptch; 1:1,000; ab53715; Abcam) and sonic hedgehog homolog (Shh; 1:500, 20697-1-AP; Proteintech) overnight at 4°C, followed by incubation with a peroxidase-coupled secondary antibody for 1 h at 37°C. The blots were visualized using an enhanced chemiluminescence kit (Amersham, UK).

### 5. Quantitative Real-Time Polymerase Chain Reaction (PCR)

Total RNA was extracted from the cells or tissue samples using the RNeasy Mini Kit (Qiagen, Valencia, CA), following the manufacturer’s instructions. Equal amounts of total RNA from each sample was prepared and reverse transcribed into complementary DNA (RR047A; Takara Bio Inc, Dalian, Liaoning, China). Supplementary Table 1 lists the specific oligonucleotide primers used. The PCR thermocycling conditions were as follows: 95°C for 30 s followed by 39 cycles of 95°C for 5 s and 56°C for 30 s using SyBr Green reagents (Biosystems, Foster City, CA). The gene expression was calculated using the 2^ΔΔCt^ method and normalized to the housekeeping gene β-actin.

### 6. Statistical analysis

All data were analyzed using SPSS version 19.0 software. Quantitative data are expressed as the mean ± SEM of at least five independent experiments for animal studies, and at least three independent experiments for each cell experimental group. The groups were analyzed using Student’s t-test and P<0.05 was considered to indicate statistically significant differences.

## Results

### 1. LXRs decreased during HSCs activation

Plating HSCs on plastic dishes can artificially cause their activation [20]. Therefore, in order to identify the underlying mechanisms of HSC activation, we tested gene expression in freshly isolated HSCs from healthy mice for 168 h. The freshly isolated quiescent HSCs had a round, phase-dense cell appearance with large numbers of refractile lipid droplets, which appeared garland-like by Oil red O staining. The purity of the HSCs was approximatively 90%, as determined by quiescent HSC marker GFAP and activated HSC marker Acta2 (Fig. 1A). Notably, collagen I and Acta2, regarded as specific markers of HSC activation, exhibited a marked upregulation during *in vitro* HSC culture (Fig. 1B). LXRα and LXRβ, and their target genes Srebp-1c and Abca1 significantly decreased during HSC activation (Fig. 1C). The protein levels of the indicators of HSC activation, LXRs and their target genes were further confirmed by western blotting. The observed trend for each protein was consistent with the corresponding mRNA expression (Fig. 1D). The results confirm that both of LXRα and LXRβ may play roles in HSCs activation.

**Figure 1:**
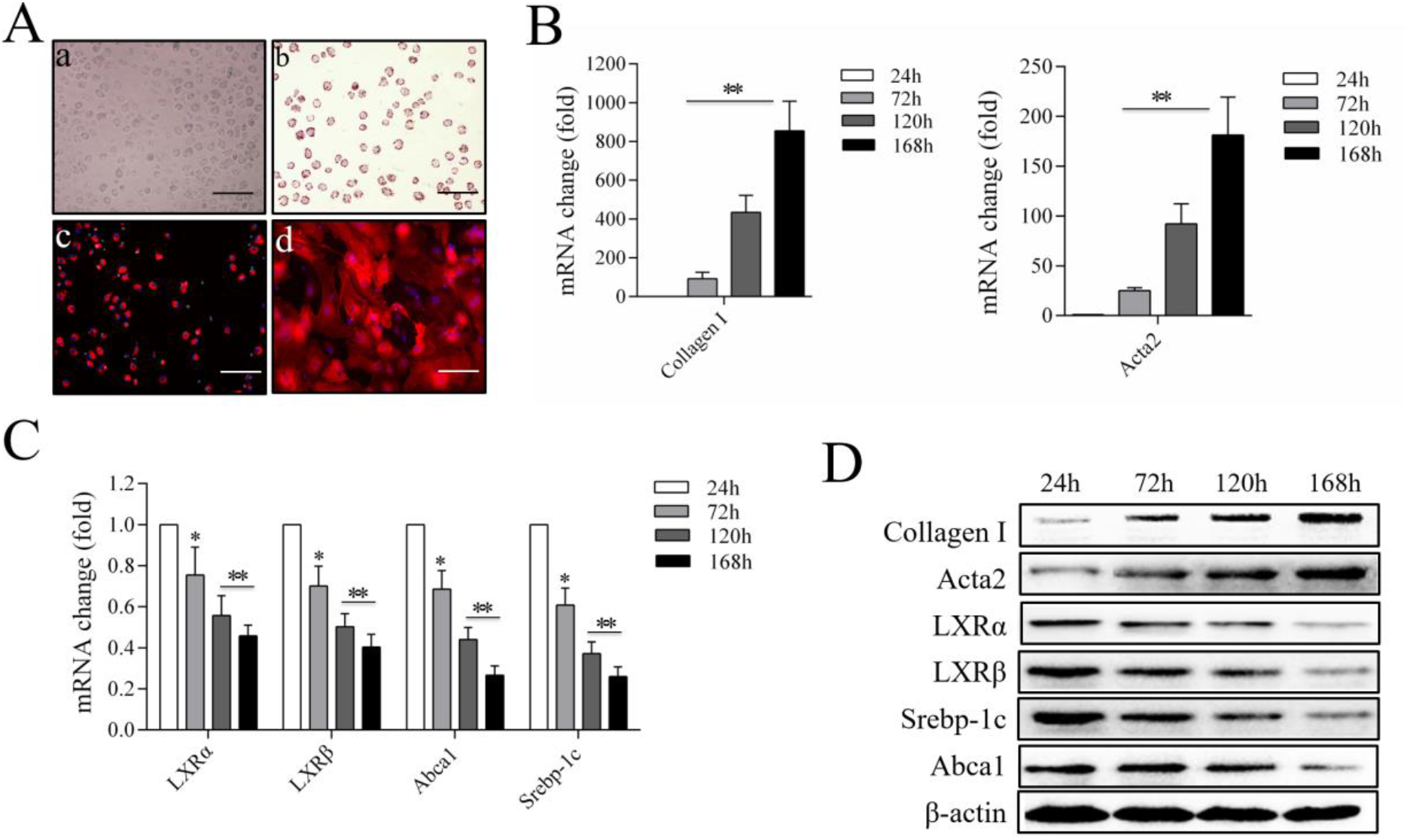
LXRs decrease during HSC activation. (A) a, Images of freshly isolated, quiescent HSCs cultured for 24 h; HSCs appear as round, phase-dense cells containing refractile lipid droplets. b, Oil red O staining of HSCs cultured for 24 h; HSCs displayed abundant garland-like lipid droplets in the cytoplasm. c, Immunohistochemical staining of HSCs cultured for 24 h; HSCs were positive for the quiescent HSC marker GFAP. d, Immunohistochemical staining of HSCs cultured for 168 h; HSCs were positive for the activated HSC marker Acta2. (B) Relative mRNA expression of collagen I and Acta2 during HSC activation. (C) Relative mRNA expression of LXRs and downstream target genes Abca1 and Srebp-1c during HSC activation. (D) Protein expression of collagen I, Acta2, LXRα, LXRβ, Srebp-1c and Abca1 during HSC activation was investigated by western blotting, using β-actin as a loading control. DAPI was used to stain the nucleus. n≥3, *P<0.05, **P<0.01. Scale bars, 50 μm.

### 2. LXRβ, but not LXRα, regulates HSCs activation in mice

According to previous studies, primary isolated HSCs are activated and induce high levels of activation markers Acta2 and collagen I *in vitro* following 72 h of culture [21]. Therefore, in the present study, freshly isolated HSCs were tested following culture for 72 h. Firstly, an LXR agonist and antagonist were tested to confirm the regulatory effect of LXRs on the activation of mouse primary HSCs. As expected, LXR agonist T0901317 led to a significant increase in the mRNA levels of Abca1 and Srebp-1c, and suppressed the activation of the mouse primary HSCs. In addition, LXR antagonist SR9238 decreased the expression of LXR target genes and notably promoted the activation of HSCs (P<0.05; Fig. 2A). Subsequently, the endogenous expression levels of LXRα and LXRβ were measured in mice primary HCs and HSCs, and normalized to the levels of the housekeeping gene β-actin. LXRα was mainly expressed in HCs, while, HSCs expressed high levels of LXRβ. And similar protein expressions were observed by western blot analysis (P<0.01; Fig. 2B). Similarly, freshly isolated HSCs exhibited strong nuclear expression of LXRβ, but almost undetectable levels of LXRα after culture for 24 h (Fig. 2C). Furthermore, we determined the effect of the knockdown of each LXR isoform on the expression of major LXR target genes and the activation of HSCs. After 72 h of culture, siRNA was used to silence LXRα or LXRβ in mouse primary HSCs. The cells in the control group were transfected with siRNA containing a scrambled sequence. Despite efficient silencing, no obvious effect by the LXRα knockdown was observed on Abca1 or Srebp-1c on the transcription level, and no significant activation of HSCs was noted (P>0.05). In contrast, LXRβ silencing led to a notable decrease in the expression levels of LXR target genes, and promoted HSCs activation, as measured by Acta2 and collagen I mRNA expression (P<0.05; Fig. 2D). The impact of LXRα and LXRβ silencing was also investigated at the protein level. Compared with the mRNA results, similar changes in the protein levels were observed by western blot analysis (Fig. 2E). Additionally, HSCs in the LXRβ-silenced group exhibited a high expression of Acta2 and displayed more fibrocyte morphological features, in contrast with the LXRα-silenced group (Fig. 2F). These results indicate that LXRβ, but not LXRα, regulates the LXR target genes and suppresses the activation of mouse primary isolated HSCs.

**Figure 2:**
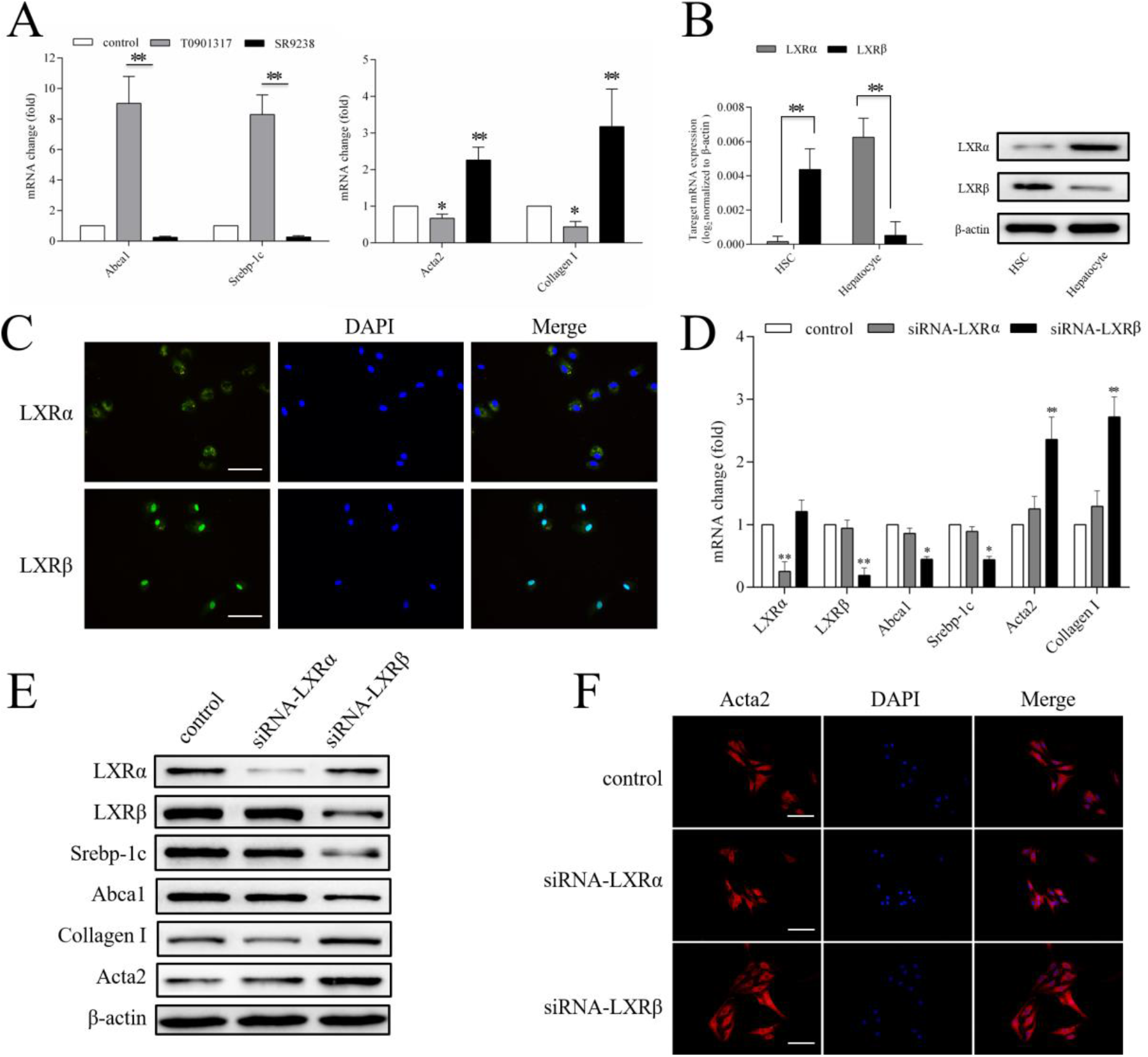
Effect of LXR knockdown on the mRNA and protein levels of LXR target genes and HSC activation. (A) Relative mRNA expression of Abca1, Srebp-1c, collagen I and Acta2. HSCs were treated with T0901317 (5 μM), SR9238 (10 μM) or DMSO for 48 h. (B) Relative expression of endogenous LXRα and LXRβ in freshly isolated HSCs. (C) Expression of endogenous LXRα and LXRβ was investigated by immunofluorescence in freshly isolated HSCs. (D) Relative mRNA expression of LXRα, LXRβ, Srebp-1c, Abca1, collagen I and Acta2. HSCs were transfected with control, LXRα or LXRβ siRNA for 48 h. (E) Protein expression of LXRα, LXRβ, Srebp-1c, Abca1, collagen I and Acta2 was investigated, using β-actin as a loading control. HSCs were transfected with control, LXRα or LXRβ siRNA for 72 h. (F) HSCs were transfected with control, LXRα or LXRβ siRNA for 72 h. Acta2 was investigated by immunofluorescence. DAPI was used to stain the nucleus. Data are presented as mean ± standard deviation. n≥3, *P<0.05, **P<0.01. Scale bars, 50 μm.

### 3. Overexpression of LXRβ inhibits CCl_4_-inducing liver fibrosis and HSCs activation in mice

CCl_4_-induced chronic liver injury was adopted to investigate whether LXRβ serves a role in HSC activation and fibrogenesis *in vivo*. After C57/BL6 mice received IP injections of CCl_4_ for 5 weeks, hepatic histopathology was evaluated by H&E, Sirius Red and Masson staining. According to the histology, the mice in the control vehicle group presented normal liver architecture, whereas the livers of the Ad-control-CCl_4_-treated mice, which received empty control adenovirus vector and CCl_4_, displayed extensive structural disorganization, with the beginnings of septa formation between adjacent vascular structures, and evidence of fibrosis completely surrounding certain parenchymal nodules. The livers of the T0901317-CCl_4_ treatment group displayed less septa formation, but were characterized by diffuse distribution of hepatic microvesicular steatosis, caused by the LXRα activation. Furthermore, the Ad-LXRβ-CCl_4_ treatment resulted in a significantly lower degree of septa formation compared with the Ad-control-CCl_4_ treatment, and overexpression of LXRβ did not induce the pathological features of liver steatosis (Fig. 3A). Subsequently, we compared the effect of T0901317-stimulated and adenovirus-mediated LXRβ overexpression on LXRs target genes at the protein level by western blot analysis. Srebp-1c and Abca1 gene expression were at low levels at basal conditions, and were drastically induced by T0901317. However, overexpression of LXRβ in the mice resulted in just a moderate increase of the expression of Abca1 and Srebp-1c, suggesting that activation of LXRβ does not significantly impact the lipid metabolism in the liver (Fig. 3B and C). Acta2 staining was performed to observe the infiltration of activated HSCs in the liver following chronic injury. The population of Acta2^+^ cells was greatly increased in the Ad-control-CCl_4_-treated mice compared with those in the mice treated with the control vehicle, indicating that chronic liver injury can produce significant HSC activation (P<0.01; Fig. 3D and E). Conversely, the number of Acta2^+^ cells was markedly lower in the LXRβ-overexpressed mice compared with that in the Ad-control-CCl_4_-treated mice (P<0.01; Fig. 3D and E). These data suggest that LXRβ suppresses the process of HSC activation and inhibits fibrogenesis in a fibrosis mouse model without resulting in liver steatosis.

**Figure 3:**
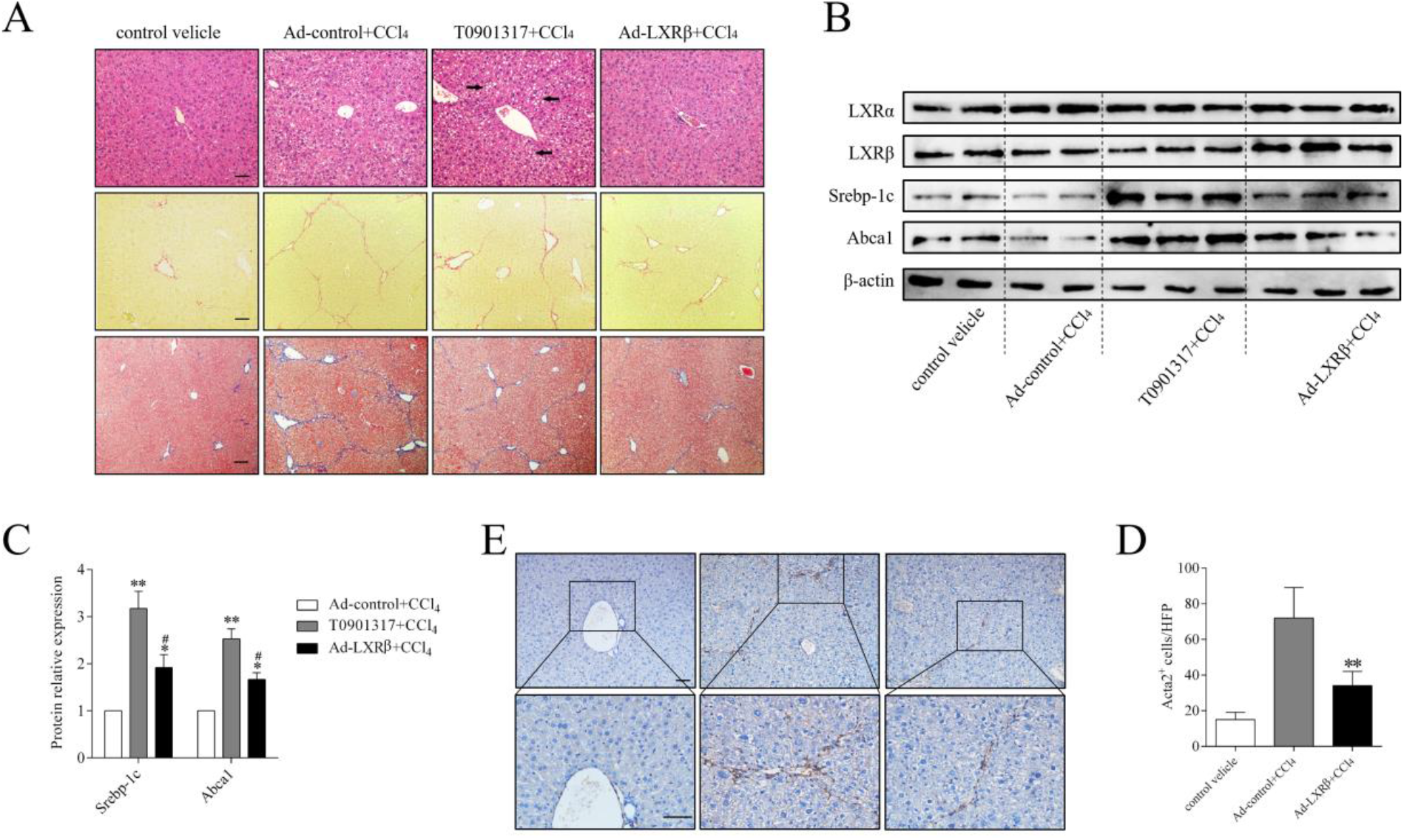
Inhibition of HSC activation in LXRβ-overexpressed mice with CCl_4>_-induced liver damage. (A) a, H&E staining of liver samples with CCl_4_-induced injury. Black arrows indicate hepatic microvesicular steatosis. b, Sirius red staining was used to evaluate the hyperplastic state of collagen, indicated with red staining. c, Masson staining was used to evaluate the fibrosis, indicated with blue staining. (B) Protein expression of LXRα, LXRβ, Srebp-1c and Abca1 was investigated by western blotting, using β-actin as a loading control. (C) Statistical analysis of the protein expression of Srebp-1c and Abca1. (D) Acta2 immunohistochemical staining of liver samples with CCl_4_-induced injury. (E) Statistical analysis of the number of Acta2^+^ cells. Data are presented as mean ± standard deviation. n=5, *P<0.05 vs Ad-control+CCl_4_ group, **P<0.01 vs Ad-control+CCl_4_ group, ^#^P<0.05 vs T0901317+CCl_4_ group. Scale bars, 50 μm.

### 4. LXRβ may inhibit HSC activation *in vivo* via Hh signaling

Hh signaling is crucial in developmental pattern formation, and stem cell growth and maintenance [22]. This signaling pathway is dormant in the livers of healthy adults, and is activated during liver injury, which triggers the production of Hh ligands and the expression of Hh target genes. It has been reported that Hh signaling is important for HSC activation and liver fibrogenesis [23,24]. As Hh signaling is negatively regulated by LXR [25,26], we hypothesized that the role of LXRβ in the regulation of HSC activation and fibrogenesis is exerted via Hh signaling. In order to test this hypothesis, the Hh pathway activity was investigated in CCl_4_-induced liver injury. Double immunofluorescence for Acta2 and Gli2, a main Hh target gene, was performed to assess Hh signaling in activated HSCs. Fig. 4A demonstrates the nuclear Gli2 localization in activated, Acta2^+^ HSCs in the livers of CCl_4_-treated mice, and the low expression in the healthy livers of the control vehicle group. Notably, the livers from the Ad-control-CCl_4_-treated group exhibited markedly increased Gli2 nuclear staining in myofibroblastic cells located in the fibrotic areas (4.6-fold increase vs. control). However, compared with Ad-control-CCl_4_-treated group, the overexpression of LXRβ significantly suppressed the number of Acta2^+^ cells as well as the nuclear expression of Gli2 (P<0.01; Fig. 4A and B). Furthermore, compared with the control vehicle group, the upregulation of Gli2 protein levels in CCl_4_-induced fibrotic liver was accompanied by an increase in collagen I and Acta2 protein levels. These results indicate that Hh signaling is involved in HSC activation and fibrosis. The overexpression of LXRβ in CCl_4_-treated mice led to a decrease in the expression of Gli2, collagen I and Acta2 (P<0.05; Fig. 4C), suggesting that LXRβ may affect Hh signaling and HSCs activation *in vivo*.

**Figure 4:**
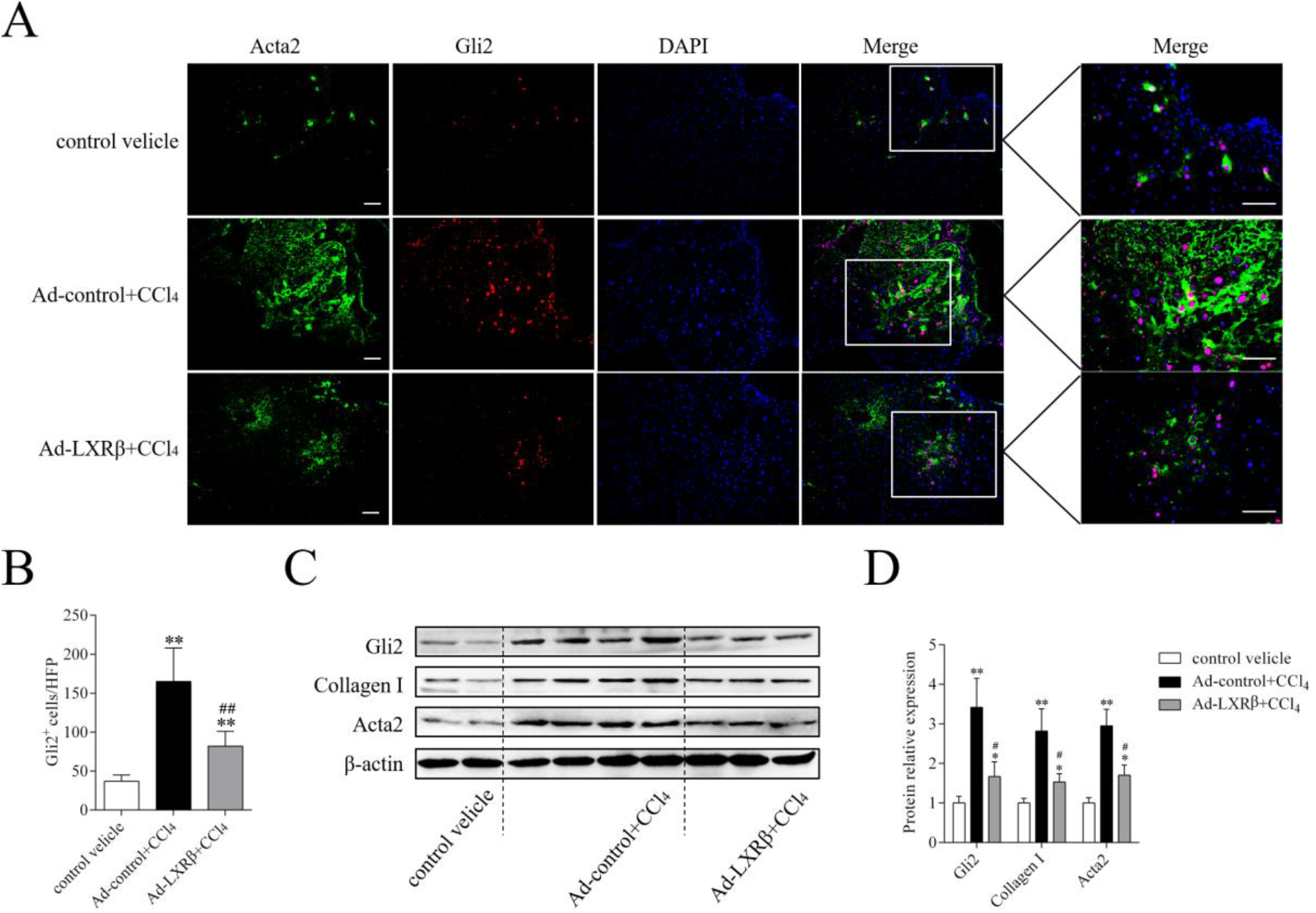
LXRβ inhibits HSC activation via Hh signaling *in vivo*. (A) Double immunofluorescence staining for the Hh target Gli2 (red), and the HSCs activation marker Acta2 (green). DAPI was used to stain the nucleus. (B) Statistical analysis of the number of Gli2^+^ cells. (C) Protein expression of Gli2, collagen I and Acta2 was investigated by western blotting, using β-actin as a loading control. (D) Statistical analysis of the protein expression of Gli2, collagen I and Acta2. n=5, *P<0.05 vs control velicle group, **P<0.01 vs control velicle group, ^#^P<0.05 vs Ad-control+CCl_4_ group. Scale bars, 50 μm.

### 5. LXRβ inhibits HSC activation via Hh signaling *in vitro*

Hh signaling activation occur at several levels of the signal transduction cascade [27,28,29]. Briefly, Hh signaling is initiated by Ptch, the cell surface receptor for the Hh ligand, binding of Shh Sonic hedgehog (Shh), a main mammalian Hh signaling ligand. This interaction permits the propagation of intracellular signals that culminate in the nuclear localization of Gli transcription factors, especially Gli2, and regulate the expression of Hh signaling target genes, including amplifications of Ptch and Gli. To further support the aforementioned hypothesis, mouse primary HSCs were isolated and cultured for 168 h. We found that the HSC activation process was accompanied by the activation of Hh signaling. Ptch and Gli2 were almost undetectable in freshly isolated, quiescent HSCs, but significantly upregulated during HSC activation (P<0.01; Fig. 5A and B). An Hh agonist and antagonist were then used to test the role of Hh signaling in the activation of HSCs. Accordingly, the mRNA expression levels of collagen I and Acta2 were upregulated by approximately 3.3- and 2.1-fold, respectively, following treatment with Hh agonist Purmorphamine compared with vehicle (P<0.01; Fig. 5C). In contrast, following treatment with Hh antagonist GDC0449, the levels of collagen I and Acta2 mRNA decreased by approximately 51 and 53%, respectively (P<0.05; Fig. 5C). These data demonstrate that Hh signaling is essential for HSC activation and fibrosis. To further confirm that Hh signaling controls HSC activation and is mediated by LXRβ, we transfected siRNA targeting LXRα or LXRβ, or adenovirus-mediated vectors overexpressing LXRα or LXRβ into HSCs, and observed their effects on Hh signaling. Silencing LXRβ in HSCs led to a marked upregulation of Hh target genes Gli2 and Ptch, and HSC activation markers collagen I and Acta2. Additionally, we observed that overexpression of LXRβ decreased the expression of Gli2 and Ptch in HSCs, and suppressed their activation, as measured by collagen I and Acta2 protein levels (P<0.05; Fig. 5D). Conversely, in the LXRα overexpression or knockdown groups, no significant differences were observed in the expression of Hh target genes or the levels of the HSC activation marker proteins, compared with the control group (P>0.05; Fig. 5D). This supports the conclusion that LXRβ is the main LXR isoform regulating HSC activation. Furthermore, double immunostaining for Gli2 and Acta2 was performed to assess the effects of LXRβ on Hh signaling in HSCs. Compared with the control vector-transfected group, the HSCs displayed higher Gli2 expression and more mesenchymal phenotype characteristics with fibroblast-like features under LXRβ knockdown conditions. On the other hand, HSCs transfected with Ad-LXRβ exhibited suppressed nuclear Gli2 expression and maintained a more quiescent phenotype (Fig. 5E). Finally, to further determine the plausible mechanisms of LXRβ inhibits Hh signaling, we transfected Ad-LXRβ in the presence of Hh agonist Purmorphamine or siRNA targeting LXRβ in the presence of Hh antagonist GDC0449 in HSCs. As shown in Fig. 5F, purmorphamine significantly induced the protein expression of Hh target genes Ptch and Gli2, and upregulated HSC activation markers collagen I and Acta2 in HSCs. Overexpression of LXRβ did not block purmorphamine-stimulated induction of Ptch and Gli2 but partially inhibited the upregulation of HSC activation markers. In contrast, GDC0449 significantly inhibited the protein levels of Ptch, Gli2, collagen I and Acta2 in HSCs, while silencing LXRβ partially rescued the inhibiting effects of GDC0449. Thus, Hh signaling is only a part of complex mechanisms for LXRβ-influenced HSC activation. Additionally, Hh ligands Shh was drastically upregulated by purmorphamine treatment, but completely abolished while overexpression of LXRβ in HSCs, indicating that LXRβ influences the Hh signaling by inhibiting Hh ligands production. Together, the results reveal that LXRβ plays an important role in inhibiting HSC activation through regulating Hh signaling.

**Figure 5:**
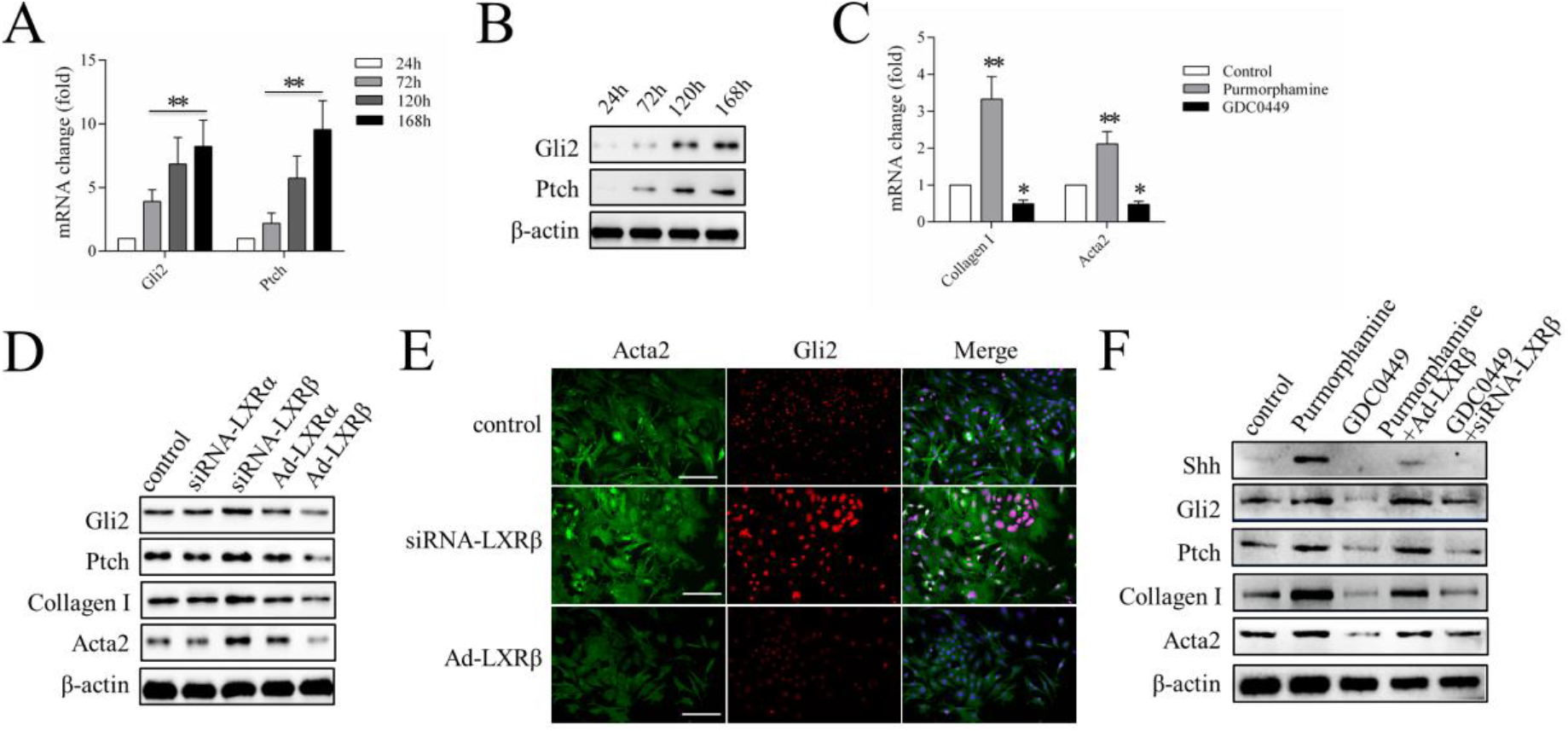
LXRβ inhibits HSC activation via Hh signaling *in vitro*. (A) Relative mRNA expression of Gli2 and Ptch during HSC activation. (B) Protein expression of Gli2 and Ptch during HSC activation, with β-actin as a loading control. (C) Relative mRNA expression of collagen I and Acta2. HSCs were treated with Purmorphamine (2 μM), GDC0449 (5 μM) or DMSO for 48 h. (D) Protein expression of Gli2, Ptch, Collagen I and Acta2 was investigated, using β-actin as a loading control. HSCs were transfected with control, LXRα or LXRβ siRNA, LXRα- or LXRβ-overexpressing adenovirus vector for 72 h. (E) Double immunofluorescence staining for the Hh target Gli2 (red) and the HSC activation marker Acta2 (green). HSCs were transfected with control, LXRβ siRNA or LXRβ-overexpressing adenovirus vector for 72 h. (F) Protein expression of Shh, Gli2, Ptch, Collagen I and Acta2 was investigated, using β-actin as a loading control. 24 h after control, LXRβ-overexpressing adenovirus vector or LXRβ siRNA transfection, HSCs were treated with Purmorphamine or GDC0449 for 72 h. n≥3, *P<0.05, **P<0.01. Scale bars, 50 μm.

## Discussion

Hepatic fibrosis cause mass mortality in the worldwide [30], but effective antifibrotic therapies are yet to be developed. Despite the fact that HSCs are a universal source for myofibroblasts in the liver, with an important function in viral, toxic, biliary and fatty liver diseases, the detailed molecular mechanisms of their activation are only partially understood, highlighting the need for further research [31]. LXRs are key regulators of hepatic lipogenesis and cholesterol homeostasis, and display anti-inflammatory and antifibrotic effects in HSCs [11,32]. Several pharmaceutical companies have been actively researching LXR agonists to activate LXRα and LXRβ, and efficiently induce the expression of LXR target genes, including Abca1 and Srebp-1c [33]. However, the activation of LXRα leading to enhanced hepatic triglyceride synthesis and adverse side effects, including liver steatosis and hypertriglyceridemia, have hindered therapeutic development. Therefore, it is important to ascertain the relative contributions of the two LXR isoforms in the process of HSC activation.

Notably, immortalized HSC lines, such as Lx2 and Hsc-T6, are poor model systems for the study of HSC activation, because these cells are already activated and resistant to transformation. For this reason, purified primary HSCs were used in the present study, which confirmed that the expression levels of the LXRs and their target genes, Srebp-1c and Abca1, were significantly decreased during HSC activation. Quiescent HSCs store large amounts of lipid and 80% of the body’s total vitamin A, but rapidly lose lipid droplets and induce the expression of activation markers when they differentiate into myofibroblasts upon stimulation. Therefore, LXRs as the key regulators of the significant changes that lipogenesis undergoes during HSC activation [12]. In contrast to HCs, in which LXRα plays a major role, HSCs mainly express high levels of LXRβ. In HSCs, LXRβ is the main regulatory isoform for the LXR target genes; knockdown of LXRβ resulted in significant decrease in Srebp-1c and Abca1 expression, and enhanced the activation of HSCs *in vitro*. On the other hand, no obvious effect of LXRα silencing was observed on the expression of LXR target genes or the activation of HSCs (Fig. 2C). Based on this result, activated LXRβ can inhibit HSC activation, while avoiding any effect on HCs, in which LXRα exerts a dominant regulatory function. Indeed, in a CCl_4_-induced liver fibrosis mouse model, overexpressed LXRβ led to the inhibition of HSC activation and fibrogenesis (Fig. 3A, D and E). Although similar findings were revealed in the T0901317-treated group, H&E staining demonstrated a diffused distribution of microvesicular steatosis in the liver (Fig. 3A). As HCs, liver sinusoidal endothelial cells and Kupffer cells, in which LXRα serves a dominant role, constitute approximately 60, 19 and 10% of the liver cell population, respectively, compared with HSCs, which mainly express LXRβ and account for just 8% [34,35,36]. In T0901317-treated mice, the two LXR isoforms were activated in parallel and led to marked upregulation of Srebp-1c and Abca1, the main regulators of triglyceride synthesis and cholesterol metabolism [37]. Overexpression of LXRβ in mice resulted in the expression of Srebp-1c and Abca1 being only moderately increased, suggesting that activation of LXRβ does not significantly affect liver lipid metabolism (Fig. 3 B and C). Therefore, the results of the present study verify the antifibrotic role of LXRs, and establish an antifibrotic mechanism for LXRβ, avoiding HSC activation-associated fibrosis and undesirable side-effect of LXRα activation simultaneously. Furthermore, we found that LXRβ regulates Hh signaling in HSC activation.

Hh signaling, a crucial developmental regulator during embryogenesis, is inactive in the healthy adult liver, but becomes reactivated during liver injury. Of the three Hh family ligands, Shh, Indian hedgehog (Ihh) and Desert hedgehog (Dhh), the Shh plays major role in liver fibrogenesis [38,39]. Under conditions of liver damage, Shh activates cell surface receptor Ptch and upregulate the expression of the main nuclear transcription factor Gli2 [40]. During this process, Hh signaling promotes the transition of quiescent HSCs to activated myofibroblastic HSCs, which produce large amounts of collagen and extracellular matrix [41]. And activated HSCs also secrete Hh ligands, which then through autocrine and paracrine role to further activate Hh signaling in a positive feedback loop. In the present study, the expression of Gli2 was low in the control group, but greatly increased in CCl_4_-treated mice, and importantly, Gli2 was mainly located in the fibrotic areas and localized in activated Acta2^+^ HSCs. According to this result, HSCs may be the main responsive cell in which Hh signaling occurs during the process of liver damage and fibrosis. Subsequently, overexpression of LXRβ significantly decreased the number of Acta2^+^ cells and their nuclear expression of Gli2 in CCl_4_-treated mice (Fig. 4A and B). Using cultured mouse primary HSCs, we verified the contribution of Hh signaling to HSC activation *in vitro* (Fig. 5A, B and C). We further confirmed the underlying molecular mechanisms between LXRβ and Hh signaling in HSC activation. Despite LXRβ suppressing Hh signaling and regulating the expression of Hh target genes Gli2 and Ptch, which are closely associated with HSC activation, no influence of LXRα on Hh signaling or HSCs activation was observed (Fig. 5D). Although it is reported that LXRα and LXRβ co-regulate several genes, and can compensate for one another in many pathological conditions, in our present study, LXRβ deletion or overexpression was not compensated by LXRα. Finally, we primarily explored the possible mechanism of HSC activation that LXRβ seems to suppress the Hh signaling by inhibiting Hh ligands production (Fig. 5F). As Hh ligand production to induce Hh signaling by at least two mechanisms, whether increasing Hh ligand expression or prolonging its half-life. Thus, much more researches need be done to explore the molecular mechanisms between LXRβ and Hh signaling.

HSCs are the most relevant source of hepatic myofibroblasts in liver fibrosis, and activated HSCs secrete several inflammatory factors that exacerbate liver injury [42]. Suppressing HSC activation and restoring the quiescent phenotype may be a promising strategy for the development of therapies against fibrosis. The results of the present study demonstrate that LXRβ regulates the activation of HSCs and prevents CCl_4_-induced fibrosis via Hh signaling, simultaneously avoiding undesirable LXRα-associated liver steatosis side-effects. Specific activators of LXRβ may be used as potential therapeutic agents against liver fibrosis.

## Conflict of interest

The authors have no potential conflicts of interest to declare.

## Acknowledgements

This study was supported by the National Natural Science Foundation of China (Grant No.82000583) and the Natural Science Foundation of Chongqing Province, China (Grant No. cstc2020jcyj-msxmX0703).

## Abbreviations

LXRα: liver X receptor α;
LXRβ: liver X receptor β;
HSCs: hepatic stellate cells;
α-SMA or Acta2: smooth muscle alpha actin;
collagen I: collagen, mainly of type I;
HCs: hepatocytes;
siRNA: small interfering RNA;
Hh: Hedgehog;
CCl_4_: carbon tetrachloride.

---

**Supplementary Figure 1:**
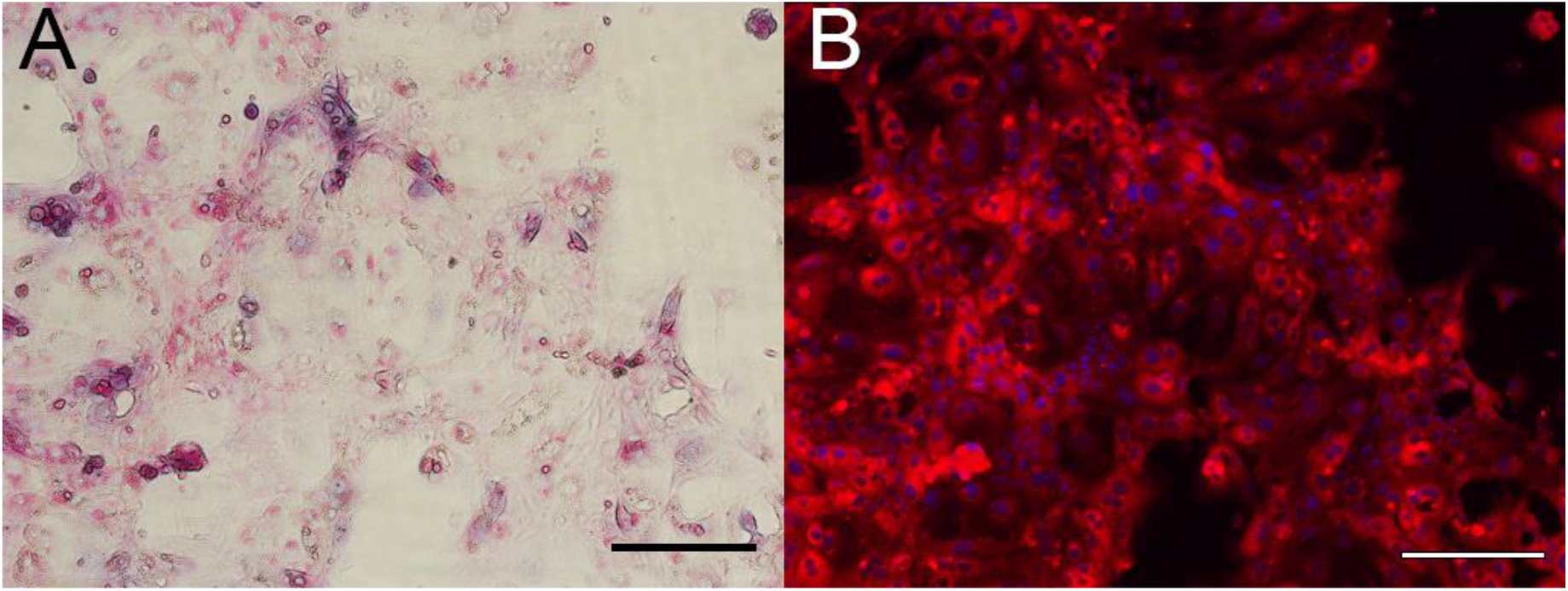
The identification of mice primary hepatocyte. A, PAS glucogen straining of freshly isolated HCs. B, Immunohistochemical staining of HCs; HCs were positive for the HC marker CK18. DAPI was used to stain the nucleus. Scale bars, 50 μm.

**Supplementary Table 1:**
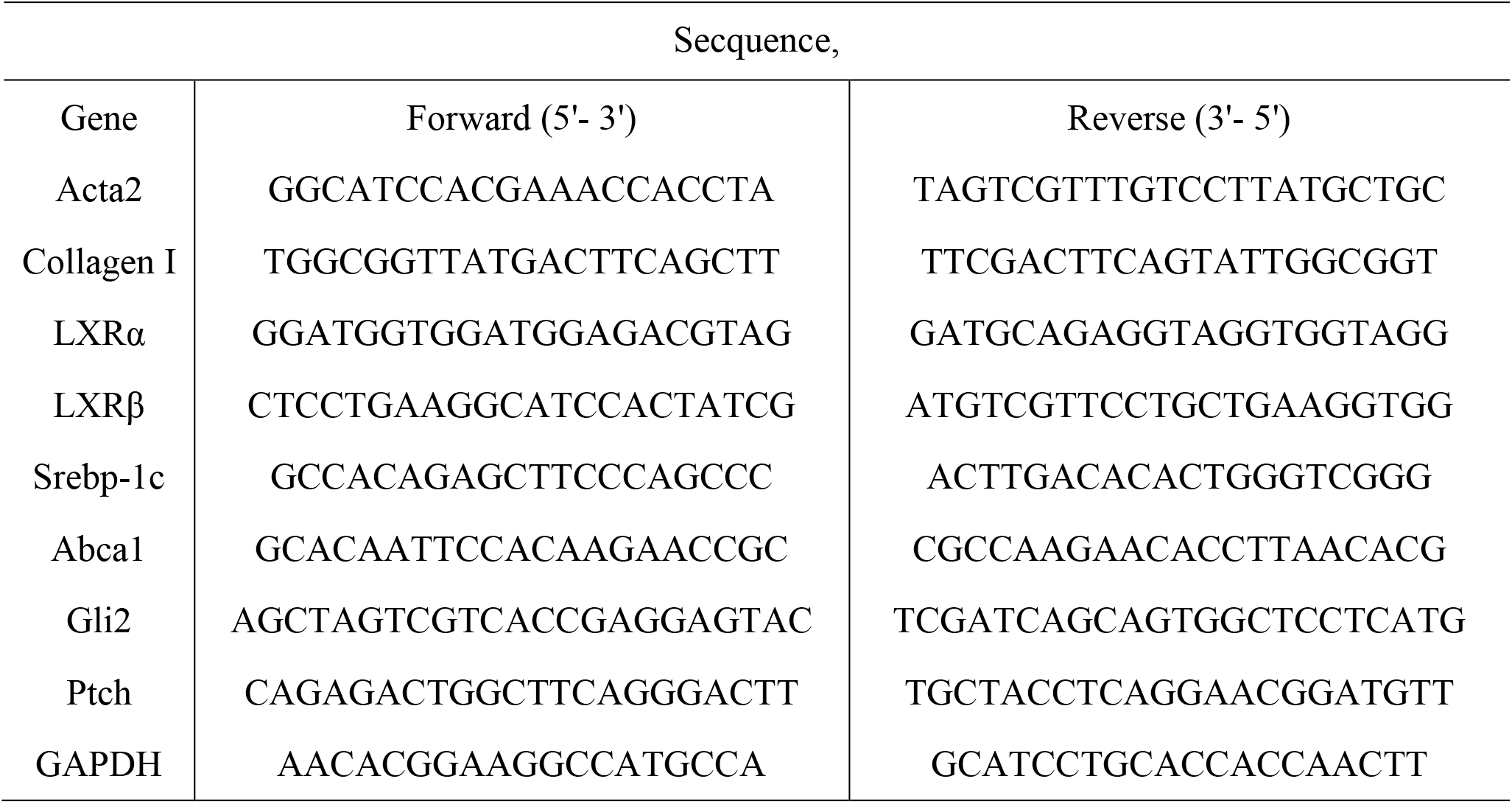
Primer and probe sequences for real-time PCR.

## Notes

Competing interests: The authors declare no conflict of interest

### Competing Interest Statement

The authors have declared no competing interest.

